# Genomic copy number predicts oesophageal cancer years before transformation

**DOI:** 10.1101/2020.02.27.967612

**Authors:** Sarah Killcoyne, Eleanor Gregson, David C. Wedge, Dan J. Woodcock, Matthew Eldridge, Rachel de la Rue, Ahmad Miremadi, Sujath Abbas, Adrienn Blasko, Wladyslaw Januszewicz, Aikaterini Varanou Jenkins, Moritz Gerstung, Rebecca C. Fitzgerald

## Abstract

Cancer arises through a process of somatic evolution and recent studies have shown that aneuploidies and driver gene mutations precede cancer diagnosis by several years to decades^1–4^ Here, we address the question whether such genomic signals can be used for early detection and pre-emptive cancer treatment. To this end we study Barrett’s oesophagus, a genomic copy number driven neoplastic precursor lesion to oesophageal adenocarcinoma^5^. We use shallow whole genome sequencing of 777 biopsies sampled from 88 patients in surveillance for Barrett’s oesophagus over a period of up to 15 years. These data show that genomic signals exist that distinguish progressive from stable disease with an AUC of 0.87 and a sensitivity of 50% even ten years prior to histopathological disease transformation. These finding are validated on two independent cohorts of 75 and 248 patients. Compared against current patient management guidelines genomic risk classification enables earlier treatment for high risk patients as well as reduction of unnecessary treatment and monitoring for patients who are unlikely to develop cancer.

---

Cancer is caused by genetic mutations that lead to an uncontrolled proliferative and invasive cellular phenotype. Mounting evidence exist that carcinogenesis occurs over many years^1^, but also that driver gene mutations can be found in physiologically normal tissues^6–10^. While early detection of cancer is one of the best strategies to improve patient survival it also poses a risk of overtreatment^11^, and accurate biomarker of cancer progression are needed. Copy number alterations are a hallmark of most cancers but are rarely found in normal tissues. These observations raise the question whether genetic signals can be decoded that identify cells at elevated risk of cancer progression.

This strategy can be tested in oesophageal adenocarcinoma (OAC) and its precursor tissue known as Barrett’s oesophagus (BE), which develops in 2%-10% of patients with chronic heartburn. OAC has a 5-year survival rate of less than 20% ^12^ and it is thus critical to develop sensitive assays for identifying individuals at risk of developing a deadly disease. However, the risk for a given patient with BE developing OAC appears to be only around 0.3% per annum and the majority of patients with BE undergo unnecessary, repeated endoscopies causing associated anxiety and health economic costs^13^.

Current surveillance programs focus on the presence and grade of dysplasia in BE patients as determined by histopathological examination of biopsies with low- and high-grade dysplasia (LGD, HGD) being used as surrogates for early cancer transformation and markers for intervention, commonly by endoscopic resection and radiofrequency ablation^14,15^. Additional risk factors for progression include increasing age, male gender, greater length of the BE segment, tobacco use, and LGD diagnosis at the initial evaluation - although these have not yet been introduced into clinical guidelines. Improvements to risk assessment have focused on identifying specific biomarkers, particularly p53 expression. Studies of p53 immunohistochemical (IHC) staining in BE biopsies suggest that including this as part of the diagnostic criteria may improve early detection programs^16–18^ as overexpression of p53 is highly specific for patients who will progress. However, p53 aberration lacks sensitivity as a predictive biomarker and is currently only recommended as an adjunct to histopathological grading in equivocal cases^19,20^.

Identification of genomic biomarkers for BE to OAC progression has been difficult, owing to the general low frequency of recurrent point mutations in either BE or OAC, perhaps with the exception of *TP53* mutations^21^. Instead, OAC is a highly copy number (CN) driven disease characterized by early and frequent genomic instability^22–26^. As ongoing genomic instability leads to a large extent of clonal diversity, multiple investigations have therefore focused on the heterogeneity of BE tissues^27^, and high levels of diversity has been associated with a higher risk of progression^28–31^.

To understand the dynamics and spatial genomic patterns of BE and their relation to OAC risk we gathered a retrospective, demographically matched, case-control cohort of patients (n=88) from a clinical surveillance program for BE. From these patients we obtained all available samples from their endoscopies (n=777), developing both a longitudinally (1-15 years per patient) and spatially (sampled along the length of the BE tissue) dense sample set. As CN alterations are the major driver of OAC we used shallow whole genome sequencing (sWGS), at an average depth of 0.4x to generate temporally and spatially resolved copy number profiles (**Methods**). Genomic risk estimates of OAC transformation were calculated using a cross-validated elastic-net regularized logistic regression model. This model was validated using an independent cohort of 76 patients (n=213 samples), alongside an orthogonal validation of the Seattle BE Study SNP array samples.

## Spatially and temporally diverse copy number profiles in Barrett’s oesophagus

Copy number (CN) patterns were examined at multiple levels of the oesophagus to understand how progressors may differ from non-progressors (**Fig. 1A**). Within a single progressor patient we find that the genomes are highly variable between each endoscopy (i.e. time) and between individual levels within an endoscopy (i.e. space). Additionally, while some patterns may be shared between individual levels, most often there is generalized disorder across the genomes.

**Figure 1:**
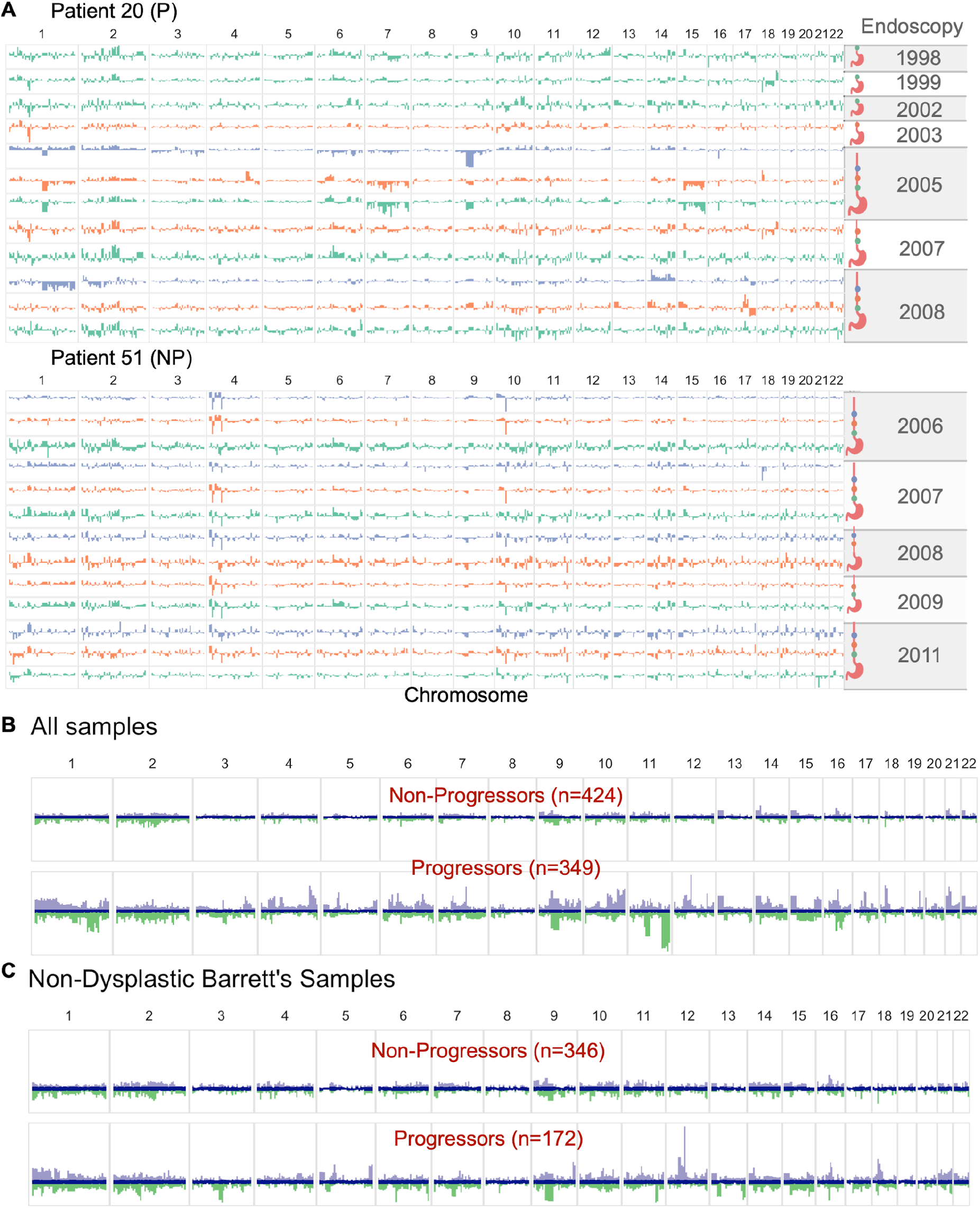
Copy number profiles in Barrett’s Oesophagous vary over space and time. Barplots showing the adjusted CN values across the genome, with relative gains shown in the positive y-axis, and relative losses shown in the negative y-axis. **A.** Genomic copy number profiles of individual samples for a progressor patient (P, top) and a non-progressor patient (NP, bottom). The colors across the chromosomes in each sample are based on the location relative to the stomach it was taken in the esophagus and the ideograms to the right of the plots show the samples that belong to a single endoscopy indicated by the year. Note the variability in the samples from the progressor patient in chromosomes 14 and 17 in contrast to the shared pattern across the non-progressor patient in those regions. **B** and **C**, distribution of relative CN values at each genomic segment across all samples in the progressor and non-progressor patient groups. The blue in the middle is the median ± 1SD, indicating a diploid genome value. Purple and green show the range of relative gains and losses, respectively. In **B** all samples regardless of pathology are plotted and a large variation in the CN between progressor and non-progressor patients is clear (i.e. chromosomes 1, 4, 9, 11). In **C** only NDBE samples from both patient groups are plotted and the progressor patients still show a much larger CN signal despite being pathologically indistinguishable.

Using the standard clinical pathology grading for dysplasia (i.e. LGD, HGD), CN changes were not driven by the degree of cytological atypia since similar profiles were observed for the non-dysplastic BE (NDBE) samples from both progressor and non-progressor patients (*n*=518; **Fig. 1B-C**). Although progressor patients have greater genome-wide CN complexity than non-progressor patients (Wilcoxon rank sum *p*=2.4×10^−6^), this information alone appeared insufficient to predict progression (**Supplementary Fig. 1A-B**).

The extensive CN data from multiple levels of the BE segment collected over many years provides an opportunity to examine the spatial architecture of BE clones as well as their temporal variation. Although shallow WGS alone does not provide the resolution to fully delineate the clonal architecture that may be present in each pooled sample, it is striking to observe higher rates of CN across large genomic regions in progressors in the early history of the disease (Wilcoxon rank sum *p*=0.026; **Supplementary Fig. 1C-D**).

## Copy-number predicts progression towards oesophageal adenocarcinoma

The CN information and a measure of overall genomic complexity from all 88 patients were used to generate a statistical classification model of progression with the endpoint HGD or intramucosal cancer (IMC; **Methods**). The probability of progression per-sample was then used to evaluate the receiver operating curve (ROC) of the classification. The cross-validated area under the curve (AUC) for this model, evaluated per sample, was 0.87 (95% CI 0.84-0.89) (**Fig. 2A**). As the model included the diagnostic samples with the most extreme CN (e.g. HGD and IMC) we evaluated a model excluding these samples (*n*=714 retained samples) which resulted in a comparable AUC of 0.93 (CI 0.91-0.95) confirming that the model was not being driven by the outliers. We subsequently evaluated a model excluding all dysplasia (e.g. LGD, HGD, IMC) to evaluate the effects and found that the classification rate was similar to the model trained on all samples (**Supplementary Fig. 9**). Aggregating predictions either per-endoscopy or per-patient did not measurably increase the prognostic accuracy, with a per-sample AUC of 0.87 (CI 0.84-0.91), suggesting that a single sample (e.g. pooled 4-quadrant biopsy) may be sufficient for prediction (**Supplementary Fig. 2**). It is important to note that this model was by design independent of demographic risk factors^32^ as our cohort was matched for sex, BE segment length, age at diagnosis, and smoking status.

**Figure 2:**
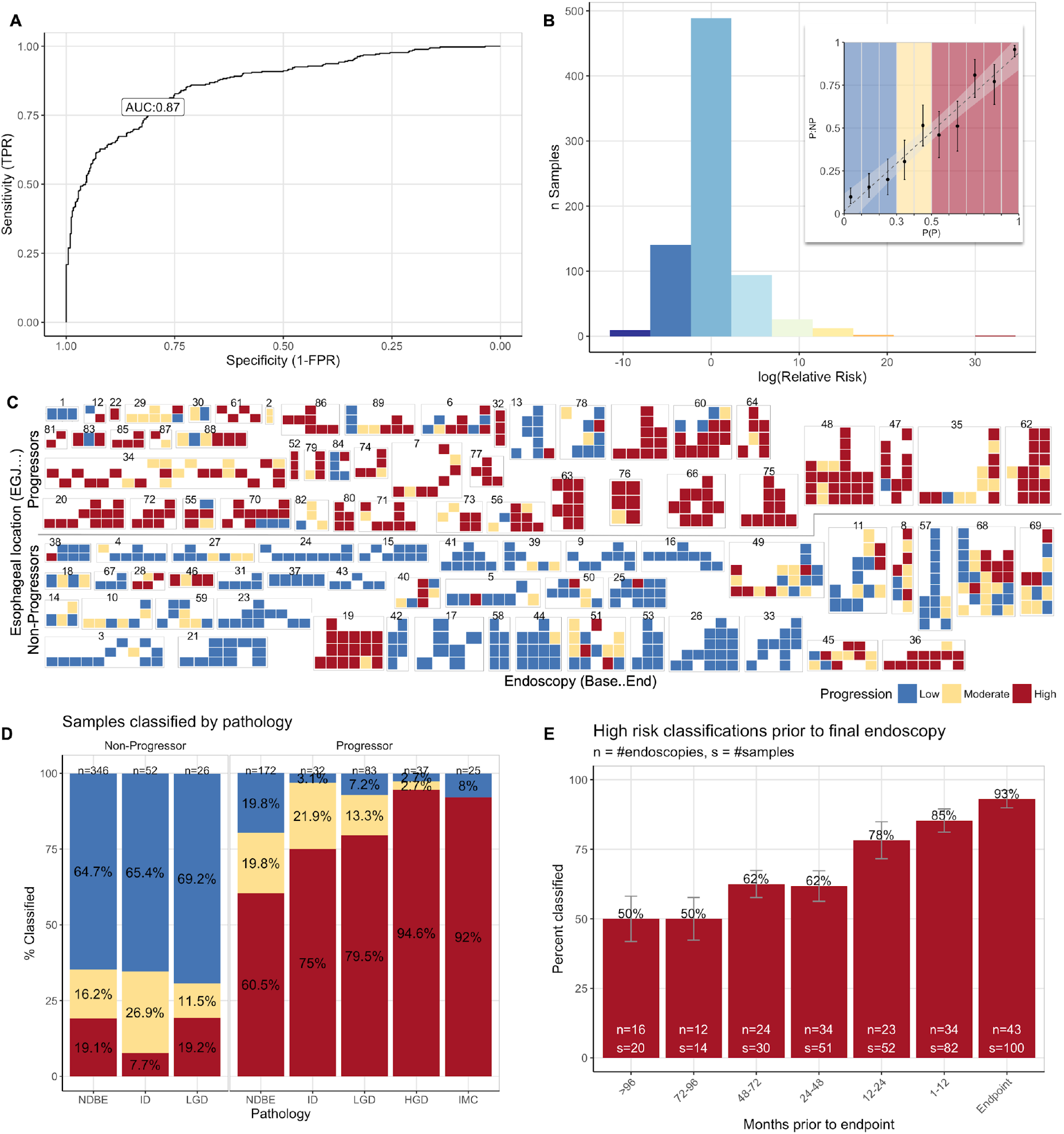
Genomic predictions of oesophageal cancer progression. **A**. ROC curve of our model predictions per-sample shows an AUC of 0.87. **B**. Histogram of the relative risk of progression across the cohort based on the leave-one-patient out predictions. The highest RR is more than 30x greater risk of progression (dark reds) while the lowest RR is at a 10x lower risk (dark blues). Inset: calibration of the predicted (x-axis) and observed (y-axis) probability of progression, evaluated in deciles. The low (blue) and red (high) risks are enriched for non-progressor and progressor patients respectively. **C**. Illustration of risk classes across all samples in the discovery cohort (n=773). The row above the line shows progressor patients, while the row below the line are non-progressor patients. Each group of tiles denotes samples from a single patient, indicated by patient number above. On the x-axis endoscopies are plotted from the baseline on the left, to the final available endoscopy on the right. The y-axis indicates the relative location of the sample starting from the esophageal-gastric junction at the bottom up the length of the BE segment. **D**. Sample risk classifications plotted per pathology (e.g. NDBE, ID, LGD, HGD, IMC). These show that our model is able to predict before pathological changes are visible in NDBE samples and **E**. that these predictions are possible up to 8 years prior to diagnosis.

## Stable predictions across space and time

Using the per-sample predictions generated by the model we evaluated the relative risk (RR) across the cohort. Those samples with the highest RR were more than 20 times more likely to progress than average, while those with the lowest RR were 10 times less likely (**Fig. 2B**). This information enabled us to calibrate our risk classes based on the enrichment of samples from progressor or non-progressor patients. The estimated probabilities were broken into three risk classes: ‘low’ (*Pr*≤0.3), ‘moderate’ (0.3<*Pr*<0.5), or ‘high’ (*Pr*≥0.5). When the resulting classes were plotted against space and time a strikingly concordant patterns emerged where most progressive patients have a higher risk of progression throughout the disease history, while non-progressive patients tend to have a very low risk of progression (**Fig. 2C**, **Supplementary Fig. 12**).

Samples from patients who progressed were classified as high risk for progression independent of histopathology (**Fig. 2D**). Most importantly, NDBE samples that belonged to progressor patients were highly likely (104/172, 60.5% of samples classified high risk) to predict progression while in non-progressor patients they are classified as low risk (224/346, 64.7% of samples). Furthermore, 50% (8/16) of endoscopies from progressor patients were classified as high risk as early as the baseline endoscopy, 8 or more years prior to transformation (classification in accordance with current diagnostic guidelines requiring only a single dysplastic sample to recommend treatment for a patient; Fig. 2E). Thus, around half of progressive BE cases harbour the CN characteristics of future dysplastic transformation early in the disease. Cases which lack early CN patterns of progression acquired these over the following years, leading to 78% (18/23) of endoscopies being classified as high risk one to two years prior to HGD/IMC diagnosis.

## Improvements over p53 IHC

One clinical tool for diagnosing progression in dysplastic samples of BE has been IHC staining for aberrant p53 expression^16^. Mutations in *TP53* are known to be early drivers of tumour progression across tumour types including OAC. As *TP53* is both clinically diagnostic and informative of the genomic development of the tumour, we performed p53 IHC staining on all biopsy levels with sufficient tissue (*n*=590, all 4-quadrants) and evaluated the predictive value of aberrant expression. We included the AUC of pathological grading for HGD or LGD as these are the basis for current surveillance guidelines.

p53 IHC had an AUC of 0.62 (CI 0.59-0.65) for progression in our cohort (e.g. HGD/IMC sample, see Methods), while diagnosis of HGD had an AUC of 0.59 (CI 0.57-0.61). In contrast, our model using genomic CN information outperformed both with an AUC of 0.87 (CI 0.84-0.89) (**Fig. 3A**). Additionally, while p53 IHC is highly specific in our dataset, it showed poor sensitivity overall. Even in progressive patients only 65% (27/41) of HGD/IMC samples were found to be aberrantly stained, dropping to 11% (13/115) in NDBE samples (**Fig. 3B**). Loss of 17p has been identified as an “early” event in the clonal evolution of OAC^20,24^. Our CN model selected −17p as one of the five most predictive features by effect size (**Fig. 3C**) However, this locus alone was not sufficient to predict progression as loss of 17p occurred in only 2% of all samples (70/773), exclusively in progressor patients where it occurred in only 20% (70/349) of samples. Of those samples, 30 were NDBE and 26 were HGD/IMC. Although this was a low proportion of overall samples it occurred in 53% of all progressor patients (24/45) and is consistent with previously reported targeted panels^33^. This loss was correlated with aberrant p53 IHC (odds ratio=10.9, *p* = 3.9×10^−11^ Fisher’s exact test).

**Figure 3:**
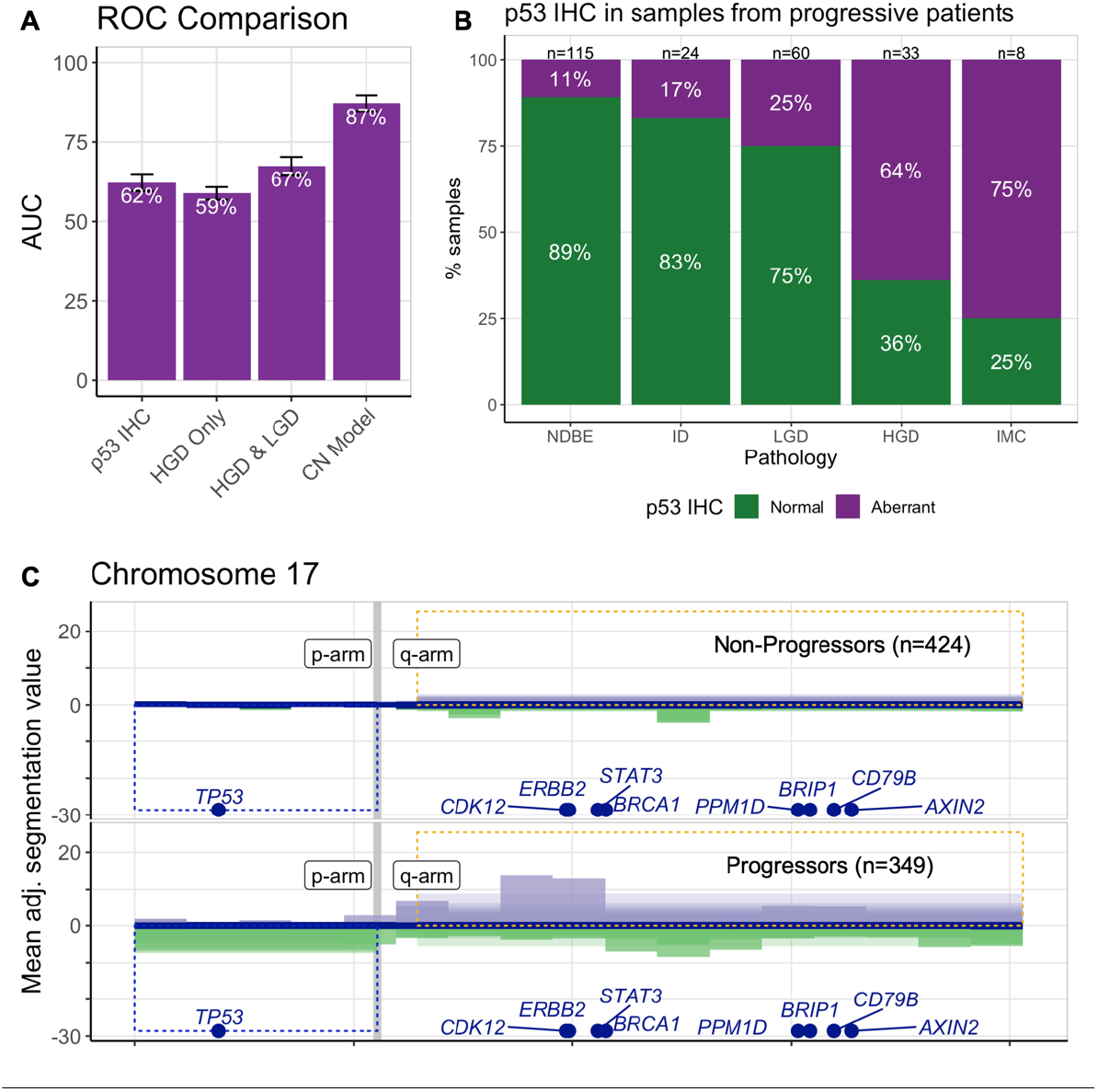
Cancer risk in relation to p53 status. **A**. Comparison of the ROC curves between predicting progression with p53 IHC, HGD, HGD & LGD, or our CN model. While the other methods have perfect (or near perfect) specificity, they have very poor sensitivity when compared to the CN model. **B**. Proportion of aberrant p53 IHC stained samples separated by pathology in samples from progressive patients. The purple bars indicate the percentage of aberrant samples for each pathology. **C**. The plot from Figure 2B zoomed in to chromosome 17 with additional bars shown for the arm-level gains (purple) or losses (green). The blue/yellow outline boxes show the genomic regions that are predictive features of the model. The blue box indicates a loss of 17p arm, while the yellow indicates gain of the 17q arm. Tumor suppressors or oncogenes are indicated at their chromosomal location at the bottom of each plot.

## Validation of model predictions

The results presented so far employed a nested leave-one out cross-validation (at the level of patients; **Supplemental Methods**). The model was further validated using an independent cohort of patients (n=75) with shallow WGS from archival samples, and SNP array data from the Seattle BE Study^34^ (n=248 patients).

An independent validation cohort of 58 patients that have not progressed and 18 patients that have progressed from NDBE to HGD or IMC was sequenced at multiple levels producing a total of 219 samples with 213 passing quality control (**Methods**). Each sample was individually evaluated using the model and classified as low, moderate, or high risk using the same criteria as in the discovery cohort. Reassuringly, the algorithm achieved an AUC of 0.81 (CI 0.74-0.87) on the validation cohort (**Fig. 4A**). 78/142 (55%) samples from non-progressor patients were classified as low risk, and 55/71 (77%) of samples from patients who progressed were classified as high risk (**Fig. 4B**). As in the discovery cohort, high risk classification of progressor patient samples was largely independent of histopathology. Additionally, this cohort was selected from available samples with no attempt to match demographics which enabled us to evaluate the extent to which clinical factors also help predict progression. However, there was no significant difference between the groups as per their progression status for sex, BE segment length, age, or smoking status (Supplemental Table 2, **Supplementary Fig. 13**).

**Figure 4:**
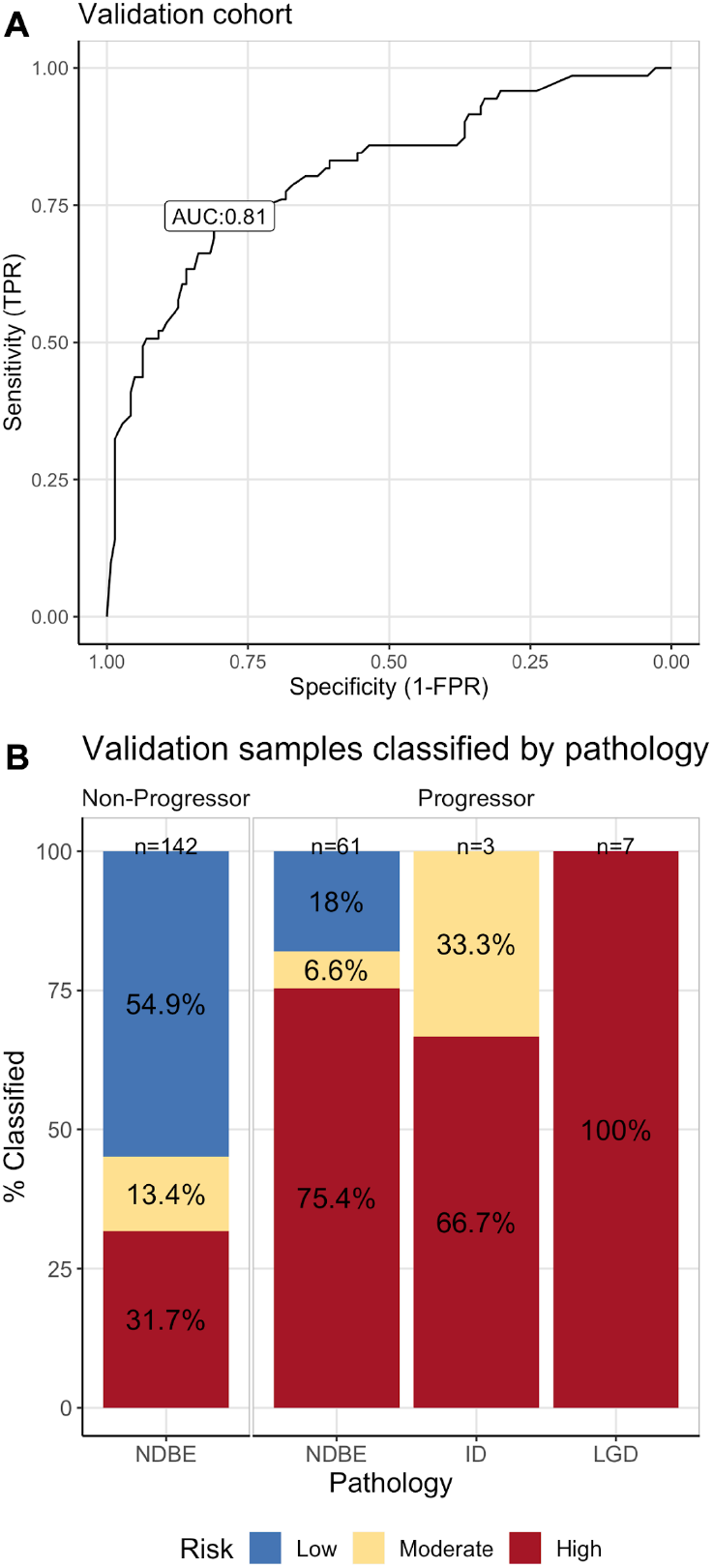
Validation of the CN prediction model on n=217 independent sWGS samples from n=76 BE cases. **A**. ROC analysis yields an AUC of 0.8 for the validation per-sample probabilities. **B.** Barplots of the risk classifications split by sample pathology in progressor and non-progressor patient groups.

Statistical algorithms can be improved by increasing the size of the dataset. To assess whether including additional patients may further improve the accuracy of CN prognosis, we conducted sub-sampling of the discovery cohort with increasing numbers of patients and model training as described before. With each increment in the number of patients the predictive accuracy increased, eventually reaching a (cross validated) AUC of 0.89 when combining all discovery and validation patients (**Supplementary Fig. 3**), indicating that a larger knowledge bank of CN and progression data from BE will continue to improve the precision of patient stratification by adding stronger statistical signal and accounting for broader biological variation.

One of the challenges in the field is having longitudinal cohorts of sufficient size to permit evaluation of predictive biomarkers. The Seattle study established in 1983 is one such cohort. Per-biopsy copy number data for the 248 patients (*n*=1,273 biopsy samples) were evaluated, although due to the historical nature of the dataset these were generated from SNP arrays rather than from sequencing. Nevertheless, predicting each endoscopic level with our model and the Seattle Study definition of progression (OAC rather than HGD/IMC; 169 non-progressors, 79 progressors) resulted in an AUC of 0.71 (95% CI 0.65-0.76) (**Supplementary Figs. 4-6**), indicating that an algorithm tailored to sequencing data may be applied to array data, but that this also introduces a certain loss of accuracy (see **Supplemental Methods and Results** for complete analysis and endpoint differences).

## Copy number stratification facilitates earlier treatment

Current guidelines for the management of BE focus on the length of the BE segment and the presence or absence of dysplasia in any biopsy sample taken during endoscopy^35,36^. Most of our patients were in surveillance prior to the current treatment recommendations for dysplasia, and hence we can compare a set of recommendations based on the current guidelines with our model using similar criteria, using our risk classifications without ignoring HGD pathology or p53 IHC where such information is available:

1. Immediate radiofrequency ablation (RFA): HGD diagnosis, or a minimum of two consecutive high-risk samples
2. Recheck 6-12 months: One high risk sample or an aberrant p53 IHC
3. Recheck endoscopy 12-24 months: One or more moderate risk samples
4. Regular surveillance 3-5 years: Two or more consecutive low risk samples

**Figure 5A-C** shows three example patients comparing current guidelines with our risk-based recommendations and describes how they differ (e.g. no RFA, fewer endoscopies). We applied these rules across our entire discovery cohort (88 patients) and evaluated the second endoscopy available in every patient to recommend based on two consecutive endoscopies.

**Figure 5:**
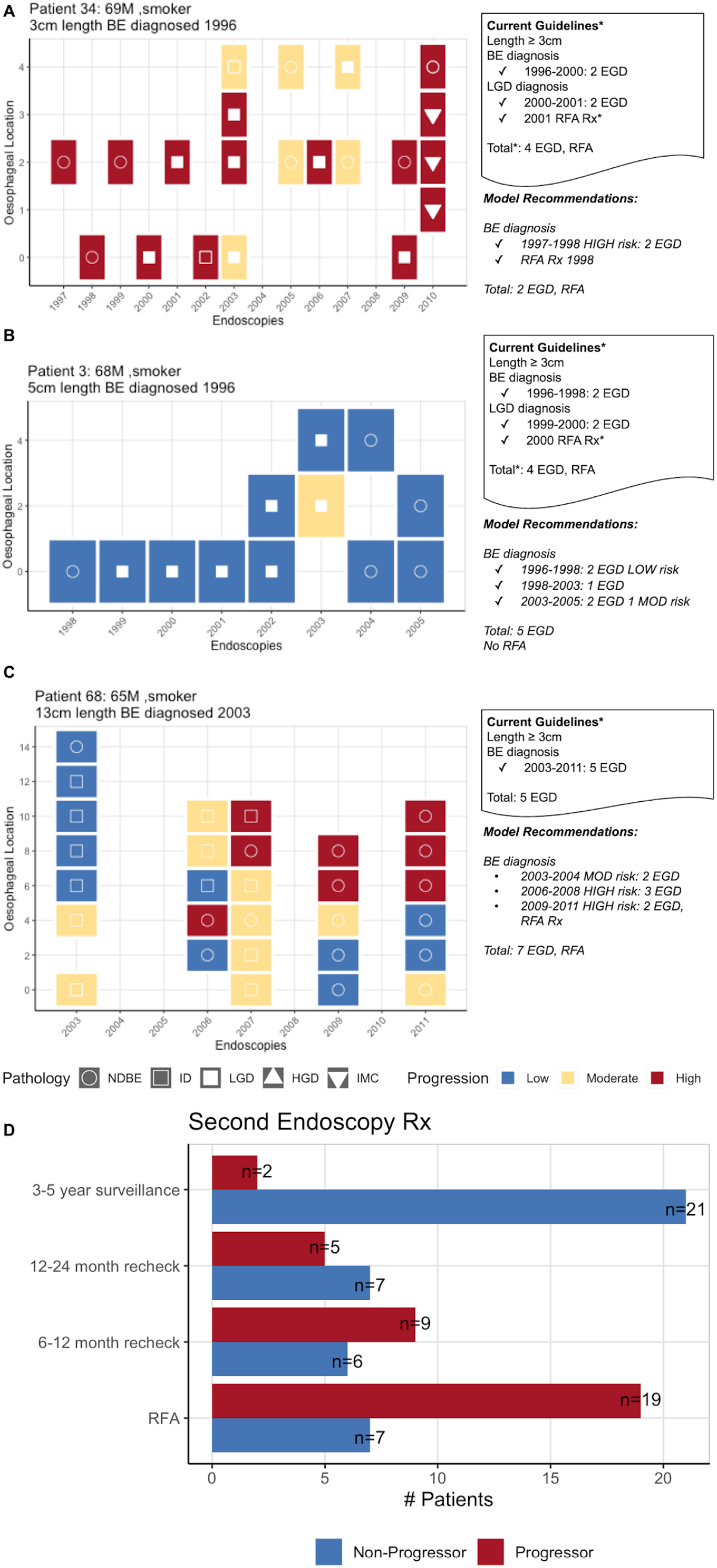
CN profiing facilitates earlier treatment and reduced monitoring. **A-C.** Individual patients with each sample plotted at the time of endoscopy. Samples are colored based on their risk class. Relevant clinical information is included above each endoscopy plot including the length of the BE segment and patient age at diagnosis. The recommendations for each patient based on the 2015 BE management guidelines are described, including the total number of endoscopies and recommendations for RFA, followed by the recommendations based on our model. **D.** Number of patients in the discovery cohort that would be recommended to follow specific surveillance or treatment protocols based on their first and second available endoscopy samples.

Using these criteria and our model predictions (**Fig. 5D**), 54% of progressor patients (19/35) would have received earlier treatment, while 51% of patients who did not progress (21/40) would have less frequent endoscopies. Only 8.5% of progressive patients (3/35) would not have been detected any sooner than their pathological diagnosis.

## Discussion

Recent evidence from the large-scale pan-cancer studies have suggested that genomic alterations are present many years before detectable disease^37^ in many cancer types. Yet it is currently unknown whether this knowledge translates to earlier cancer detection and prevention due to lack of accessible pre-malignant tissues and longitudinal observations. BE constitutes a known pre-malignant condition with historical follow-up to test whether genomic medicine can contribute to early cancer detection. Previous studies of BE progression have shown that genomic changes are present prior to cancer progression and differ in patients who do ultimately develop cancer. A number of clonal and genetic diversity measures have been proposed to stratify patients, including copy-number losses or copy neutral loss of heterozygosity^28,30,38^. Single-cell studies have further supported these diversity measures, and suggested that the higher the clonal diversity the earlier BE tissues are set to progress^29^.

Moreover, our analysis has shown that this depth of information, requiring complex sample preparations and expensive workflows often down to a single cell level, may not be needed to measure the risk of progression. While copy number profiles were indeed highly variable across space and time, in accordance with ongoing chromosomal instability, these translated into surprisingly stable predictions of a patient’s risk of progression. Further, these single-sample predictions (i.e. from all biopsy quadrants at one level) were almost as accurate as aggregated data from multiple biopsies across the entire endoscopy (usually 8-20 samples), showing that despite high levels of divergence there are common patterns of copy number alterations indicative of progressive disease. Furthermore, the observation that 50% of cases could be identified ten years prior to histopathological signs of cancer progression emerged shows that predisposition for malignancy is encoded early in the genomes of cancer precursors. That the other 50% appear to develop more slowly could be seen due to the rich longitudinal sampling of this cohort and provides insight into the evolutionary development of different clonal populations. Importantly, CN analyses based on individual biopsy samples could be conducted as part of standard monitoring practice.

Perhaps most interestingly for biomarker investigations is that, while our statistical model selects some genomic regions of instability as features that are known to be early drivers of OAC (e.g. *TP53*), few other features have any clearly associated tumour suppressor genes or other cancer-related activity (**Supplementary Table 3**). The heterogeneous nature of BE would partly explain the differences between the features our model selects as significantly contributing to progression from those found in previous studies^30^, however, there is currently no clear functional explanation for most of the features identified. Perhaps, as with other copy-number driven cancers, it is the sum of many small changes and the breakdown of gene regulatory control that fuels oncogenicity.

Ultimately, the combined use of genomic technologies, clinical samples and statistical modelling presented here is an example of how genomic medicine can be implemented for early detection for cancer. As the calculations show, genomic risk stratification has a realistic potential to enable earlier intervention for high-risk conditions and at the same time reduce the intensity of monitoring or even reduce over treatment in cases of stable disease. Whether a similar potential can be realised in other cancers depends on the accessibility of tissues and also the availability and invasiveness of treatments. Blood samples can be relatively easily obtained enabling monitoring conditions such as clonal haematopoiesis and trialling of novel treatments. For some lesions that are comparatively easy to treat or excise, such as neoplasms of the skin or colonic polyps, there may be less need for additional genomic stratification. Yet for many other cancers with existing screening programs including breast, prostate and lung cancer, sequencing of suspected and confirmed premalignant samples from longitudinal cohorts will provide answers whether a genomic basis for refined early detection and preventative treatments exists.

## Methods

### Patient cohorts

A nested case-control cohort of 90 patients were initially recruited to this study from patients that had been under surveillance for BE in the East of England from 2001 to 2016 for a total of 632 person years. Permission to analyse existing clinical diagnostic samples was approved by the relevant institutional ethics committees (REC 14-NW-0252). Cases comprised 45 patients who progressed from NDBE to HGD or IMC with a minimum follow-up of 1 year (mean 4.6 ± 3.7 years). Controls were 45 patients who had not progressed beyond LGD with a minimum follow-up of 3 years (mean 6.7 ± 3.2 years). Cases and controls were matched for age, gender, and length of BE segment (**Supplementary Table 1**). Patients had endoscopies at intervals determined by clinical guidelines with 4-quadrant biopsies taken every 2cm of BE length (Seattle protocol). One non-progressor patient revoked consent prior to analysis, and a second non-progressor was later removed during analysis when multiple comorbidities affecting the oesophagus were identified.

Each sample was graded by multiple expert GI histopathologists using current clinical guidelines for IMC, HGD, LGD, indeterminate (ID), NDBE. A single biopsy graded as HGD or IMC was considered the endpoint for progression as patients were immediately recommended treatment in the clinic. Since 2014 patients with LGD are also routinely treated with RFA making analysis of the real rate of progression difficult.

An independent unmatched cohort of 75 patients was subsequently selected from clinical surveillance patients for model validation. This cohort was comprised of 18 patients who had progressed from NDBE to HGD or IMC with a minimum follow-up of 1 year (mean 6.1 ± 3.4 years) and 58 patients who had not with a minimum follow-up of 1.5 years (mean 5.4 ± 3.0 years). The earliest available endoscopy samples subsequent to BE diagnosis were obtained to assess future risk. No endpoint samples (e.g. HGD or IMC) were included. (**Supplementary Table 2**).

All UK patients had previously given informed consent to be part of the following studies: Progressor study (REC -10/H0305/52), Barrett’s Biomarker Study (REC -01/149), OCCAMs (REC 07/H0305/52 and 10/H0305/1), BEST (REC 06/Q0108/272) BEST2 (REC 10/H0308/71), Barrett’s Gene Study (REC 02/2/57), Time& TIME 2 (REC 09/H0308/118), NOSE study (REC 08/H0308/272), Sponge study (REC 03/306).

Patient samples from the Seattle Barrett’s Esophagus Study^34^, which use SNP arrays as an orthogonal measure of CN with an endpoint of OAC, were also included for further validation (**Supplemental Methods and Results**).

### Tissue Sample Processing

Formalin fixed, paraffin embedded (FFPE) tissue samples gathered during routine surveillance endoscopies were processed from scrolls, without microdissection. Following the Seattle protocol for endoscopic surveillance 4-quadrant biopsies were taken every 1-2cm of the Barrett’s length at each endoscopy per patient. At each 1-2cm length the quadrant biopsies were pooled for sequencing as a single sample to ensure sufficient DNA (75ng) was present.

Patient samples from the discovery (*n*=88) cohort with sufficient tissue (*n*=590) were also immunohistochemically stained (IHC) for p53 and graded by an expert pathologist as aberrant or normal.

### Shallow whole genome sequencing pipeline

Single-end 50-base pair sequencing was performed at a depth of 0.4X on the Illumina HiSeq platform. Sequence alignment was performed with BWA^39^ v.0.7.15, and pre-processing of the reads for mappability, GC content, and filtering was performed with QDNAseq^40^ in 50kb bins. Single-end 50-base pair sequencing was performed at a depth of 0.4X on the Illumina HiSeq platform. Sequence alignment was performed with BWA^39^ v.0.7.15, and pre-processing of the reads for mappability, GC content, and filtering was performed with QDNAseq^40^ in 50kb bins.

Samples were jointly segmented per patient using the piecewise constant fit function (pcf) in the R Bioconductor copynumber v1.16 package^41^. Input to this function was the GC adjusted read counts from QDNAseq (Supplementary Figs. 7-8). Per-segment residuals were calculated and the overall variance across the median absolute deviation of the segment residuals was derived as a per-sample quality control measure. Samples with a variance greater than 0.008 were excluded from analysis. Samples were jointly segmented per patient using the piecewise constant fit function (pcf) in the R Bioconductor copynumber v1.16 package^41^. Input to this function was the GC adjusted read counts from QDNAseq. Per-segment residuals were calculated and the overall variance across the median absolute deviation of the segment residuals was derived as a per-sample quality control measure. Samples with a variance greater than 0.008 were excluded from analysis.

### Statistical methods

We encoded all of the sWGS copy number data on a genome-wide scale by taking a weighted average of the segmented values per 5Mb windows, and mean standardizing per genomic window. In order to evaluate chromosomal instability on a larger scale we averaged the segmented values across chromosome arms and adjusted the each 5Mb window by the difference between the window and the arm. The resulting data was 589 5Mb regions and 44 chromosome arms. We additionally included a measure of genomic complexity by summing, per-sample, the 5Mb regions that had copy-number values two standard deviations from the mean.

We performed elastic-net regression was with R glmnet^42^ package to fit regression models with varying regularization parameters. 5-fold cross validation repeated 10 times was performed on a per-patient basis removing all samples from 20% of patients in each fold. This process was performed in three conditions: using all samples; excluding HGD/IMC samples; excluding LGD/HGD/IMC. The two exclusion conditions were performed in order to assess the contribution of dysplasia to the classification rate of the model (**Supplementary Fig. 9**).

The model was additionally tuned on two parameters: 1) QDNAseq bin size; and 2) elastic-net regression penalty, between 0 (ridge) and 1 (lasso). We assessed the cross-validation classification performance of the model at multiple QDNAseq bin sizes, and at multiple regression penalties. We selected the final QDNAseq bin size by comparing the leave-one-patient out predictions from the discovery cohort, to the model predictions for the validation. This was done to minimize the batch errors in the raw data (**Supplementary Figs. 10-11**). For the regression penalty parameter, all of the models had a cross-validation classification rate of 72-75%. We therefore selected the parameter that limited the number of coefficients (n=74) and was not full lasso (e.g. 0.9). Coefficients determining the logarithmic relative risk change stemming from a unit change were calculated for each genomic region selected.

Subsequently, a leave-one out analysis (excluding all samples of an individual) was performed to generate predictions for all samples from a single individual and estimate the overall prediction accuracy using the area under the receiver operating characteristic curve using the R pROC^43^ package. Confidence intervals reported denote the 95% range.

## Data availability

Sequencing data and associated metadata that support this study have been deposited in the European Genome-phenome Archive under accession (in process; link added upon publication). The code and model that support these findings have been provided as an R package in a GitHub repository (https://github.com/gerstung-lab/BarrettsProgressionRisk).

## Author contributions

S.K. developed the statistical methods, analysed data, and wrote the manuscript and supporting information with input from E.G. and M.G. E.G. put together the discovery cohort, developed the sWGS methods, generated the sWGS data, and curated the clinical information with support from A.V.J. S.K. and E.G. are joint first authors. The initial processing pipeline was developed by D.C.W., D.J.W. and M.E., and provided input to the data analysis for the sWGS data. W.J., R.R. and A.M. identified, collected, and assessed pathology for patient samples. S.A. and A.B. sequenced the validation cohort samples. R.C.F. initiated and jointly supervised the study. R.C.F. and M.G. wrote the manuscript and are joint corresponding authors.

## Acknowledgements

We thank the patients who donated tissue samples to this project. The laboratory of R.C.F. is funded by a Core Programme Grant from the Medical Research Council (RG84369). This work was also funded by a United European Gastroenterology Research Prize (RG76026). We thank the Human Research Tissue Bank, which is supported by the UK National Institute for Health Research (NIHR) Cambridge Biomedical Research Centre, from Addenbrooke’s Hospital. Additional infrastructure support was provided from the Cancer Research UK-funded Experimental Cancer Medicine Centre. We also thank Brian J. Reid, Patricia C. Galipeau, and Carissa A. Sanchez from the Fred Hutchinson Cancer Research Center for their time and help in understanding their data, as well as Alexander Wolfgang Jung from the EBI for his time.

