## Supplemental figures and tables for "Genomic copy number predicts oesophageal cancer years before transformation"

### Supplementary Information

#### Table of Contents

*Supplementary Methods & Results* 1

*Seattle Barrett's Esophagus Study Cohort* 2

**Seattle Patient Consent** 2

*Supplementary Methods* 2

**Seattle Barrett's Esophagus Study Cohort** 2

Pathology & Demographics 2

SNP Array Pipeline 2

*Supplementary Results* 3

**Seattle Barrett's Esophagus Study Cohort** 3

*References* 5

*Supplementary Tables* 6

*Supplementary Figure Legends* 11

### Seattle Barrett's Esophagus Study Cohort

All 248 patients (n=1272 biopsies) with Barrett's esophagus without EAC at either the baseline or penultimate endoscopy were included from the case-control study in the Seattle Barrett's cohort[1]. 79 patients who progressed to EAC and 169 patients who did not progress. Importantly, non-progressive patients in the Seattle Study could have HGD as progression was only defined as EAC. Clinical endoscopy biopsies were obtained following the Seattle protocol for endoscopic surveillance. One fresh-frozen research biopsy per 2cm length of BE segment was obtained from flat mucosa without visible IMC, and BE epithelium was isolated for SNP arrays. Further details can be found in the original publication.

#### Seattle Patient Consent

Seattle Barrett's Esophagus Study research participants contributing clinical data and biospecimens provided written informed consent, subject to oversight by the Fred Hutchinson Cancer Research Center IRB Committee D (Reg ID 5619).

### Supplementary Methods

#### Seattle Barrett's Esophagus Study Cohort

##### Pathology & Demographics

169 patients did not progress to EAC from their initial diagnosis, while 79 patients did progress. Patients and samples were not matched, length of follow-up for cases may have been as short as 6 months. Further details can be found in the original publication.

##### SNP Array Pipeline

DNA from epithelial purified fresh-frozen samples were processed on HumanOmni1-Quad v0.1 bead arrays. Single-sample copy number calls, as well as purity and ploidy estimates were obtained using ASCAT without matched germline. The original estimate of the percentage of the genome that had been altered was obtained from the initial analysis reported by Li et.al. Data was encoded for model prediction on a per-endoscopy (e.g. timepoint) basis, merging all biopsies per endoscopy as per the processing pipeline described above (e.g. weighted average of CN values per 5Mb window), for both 5Mb segments and 44 chromosomal arms. This resulted in 490 merged samples across a total of 634 genomic locations. Each sample was mean-adjusted for diploid state and z-normalized using the sWGS mean and standard deviation per-genomic window. Each timepoint sample was then predicted using the sWGS trained model described.

### Supplementary Results

#### Seattle Barrett's Esophagus Study Cohort

We evaluated predictions using our model in the processed SNP data using the area under the curve (AUC) from a receiver operating curve (ROC) analysis. Using all BE merged timepoint samples with progression status as stated in the original study the AUC is 0.71 (95% CI 0.65-0.76). (Supplementary Fig. 4A).

Risk classification was simplified for the SNP cohort to 'low' ($P< 0.5$) and 'high' ($P\geq0.5$). This is equivalent in the sWGS data to merging all 'moderate' class samples into the 'low' class and is the measurement used to assess the initial training classification of the model. Using the low and high classes we evaluated each timepoint by the pathology grade provided. Overall, the timepoints from patients who progressed to EAC were classified as 'high' risk (61.5-88.9%) regardless of the specific pathology. HGD and EAC timepoint samples in progressor patients were classified as 'high' risk 73.3-87.5% respectively. More importantly, the timepoints with NDBE pathology in the progressive patients were classified as 'high' risk in 71.4%. In non-progressor patients our model predicted a higher rate of false positives in

NDBE, ID, and LGD but overall 55-61.3% of timepoint samples in each of these groups were classified as ‘low’ risk (Supplementary Fig. 4B). This is consistent with the UK validation cohort pathology risk classifications as well.

In the Seattle cohort patients with HGD were not considered to have progressed, and 36 timepoint samples (from 26 patients) fall into this group. Interestingly, 72.2% of these timepoints were classified as ‘high’ risk. As the sWGS model was trained to an endpoint of HGD or IMC this suggests that overall genomic complexity increases in HGD and that these patients are at a higher overall risk of progression as has been previously noted[2,3]. This was not the case for LGD in any of the three cohorts.

As the AUC in the Seattle data was much lower than the sWGS discovery or validation data we further explored the differences in the data. Overall SNP data provides more specific copy number information than the sWGS data can. However, the sWGS model relies on many genomic regions with very small effect sizes as the technology is genome-wide rather than targeted, and for this the SNP data provides less information overall. We can see this by training a model using the same methods used in sWGS (Supplementary Fig. 5). The result is that the number of stable coefficients used for prediction (27) is less than half that of the model we trained on sWGS data (75) and of those only seven are shared between them including the complexity score (cx) and 17p arm.

We subsequently filtered the SNP timepoint samples to select those where the mean percentage of the genome altered (per ASCAT) for biopsies per timepoint was greater than 1% and where ASCAT assessed their purity at less than 95%. These cutoffs were selected by comparing the ASCAT blood/gastric normal sample calls to all BE biopsy samples (not the merged timepoints). As the CN calls in SNP data are more sensitive overall than in sWGS our model is unlikely to identify relative CN changes in genomes that are essentially diploid.

After excluding these low CN samples and non-progressor samples that were whole-genome duplicated (a hallmark of EAC), we analyzed 286 timepoints across 186 patients (117 non-progressors, 69 progressors). In this group the AUC increases to 0.76 overall. We further assessed the predictive accuracy of our model with respect to time. In the baseline timepoint (T1) our model results in an AUC of 0.72, while in the penultimate timepoint the AUC increases to 0.82 (Supplementary Fig. 6). This is consistent with our sWGS data where we are able to identify early samples as high risk, but at a slightly lower rate than samples closer in time to clinical diagnosis.

Overall the AUC in these data are lower than those on the sWGS samples. Some of this difference can be explained due to the different sampling: in the Seattle study each 1-2cm of the BE segment had only a single biopsy while each UK sample was a pool of the 4-quadrant biopsies. Despite these differences, the AUC do show that the model works across different data platforms, and the overall findings with respect to pathology and time are consistent.

#### Supplementary Tables

| UK Discovery Cohort | Non-Progressors | Progressors | P-value (Test) |
|---|---|---|---|
| Total Patients | 43 | 45 | |
| Total Samples *(passed QC)* | 424 (424) | 353 (349) | |
| Range years follow-up (Mean ± SD) | 3-13 (6.7 ± 3.2) | 1-15 (4.6 ± 3.7) | |
| Mean age at diagnosis (years) | 59.7 ± 10 | 62.4 ± 9.7 | P=0.4 (Wilcoxon rank-sum) |
| Mean Barrett's segment length in cm *(not recorded)* | 1.3 ± 2.5 *(3)* | 1.4 ± 2.1 *(15)* | P=0.7 (Wilcoxon rank-sum) |
| Gender Female:Male | 11:32 | 8:37 | P=0.4 (Fisher's exact) |
| Smoking Yes:No *(not recorded)* | 16:6 *(21)* | 19:5 *(21)* | P=0.7 (Fisher's exact) |
| Samples IHC stained for p53 *(Not stained)* | 347 *(77)* | 243 (110) | |
| Mean number of endoscopies ± SD | 5.1 ± 1.8 | 4.2 ± 3.2 | |
| Reported pathology - number of samples | | | |
| Non-dysplastic Barrett's (NDBE) | 346 | 176 | |
| Indeterminate (ID) | 52 | 32 | |
| Low-Grade Dysplasia (LGD) | 26 | 83 | |
| High-Grade Dysplasia (HGD) | N/A | 37 | |
| Intramucosal Carcinoma (IMC) | N/A | 25 | |

**Supplementary Table 1: UK Discovery Cohort Demographics**

Patients were matched for age at diagnosis, BE segment length and gender. Patients who did not progress to HGD/IMC had a minimum 3 years follow-up, patients who progressed to HGD/IMC from NDBE had to have a minimum 1 year follow-up.

| **UK Validation Cohort** | **Non-progressors** | **Progressors** | **P-value (Test)** |
|---|---|---|---|
| Total Patients | **58** | **18** | |
| Total Samples *(passed QC)* | **143 *(142)*** | **76 *(71)*** | |
| Years follow-up (Mean ± SD) | **1.5-12.3 (5.4 ± 3.0)** | **1.6-13 (6.1 ± 3.4)** | |
| Mean age at diagnosis (years) | **67.1 ± 13** | **62.8 ± 9.7** | **P=0.09 (Wilcoxon rank-sum)** |
| Gender Female:Male *(unknown)* | **14:42** | **4:14** | **P=1 (Fisher's exact)** |
| Mean Barrett's segment length in cm | **6.3 ± 3.1** | **5.4 ± 2.9** | **P=0.22 (Wilcoxon rank-sum)** |
| Smoking Yes:No *(not recorded)* | **14:16 *(27)*** | **11:4 *(3)*** | **P=0.1 (Fisher's exact)** |
| **Reported pathology – number of samples** | | | |
| Non-dysplastic Barrett's (NDBE) | **147** | **63** | |
| Indeterminate (ID) | **0** | **3** | |
| Low-Grade Dysplasia (LGD) | **0** | **10** | |

**Supplementary Table 2: UK Validation Cohort Demographics**

In the validation cohort patients were not matched for gender, follow-up, or BE segment length. In this group we aimed to evaluate a single timepoint from the earliest possible full surveillance endoscopy. In the progressor patients we did have a couple more timepoints as we were not certain that we would have enough DNA.

| Coefficient | CV(RR) | Gain/Loss |
|---|---|---|
| 11:106076549-110898209 | 21.083 | Loss |
| 15:43942026-48824472 | 20.042 | Loss |
| 1:189430472-194415484 | 15.924 | Loss |
| 17:1-22200000 | 8.546 | Loss |
| 6:24445010-29334011 | 5.814 | Gain |
| 17:25800000-81195210 | 5.090 | Gain |
| 7:59676999-64650081 | 2.785 | Loss |
| 16:19022054-23777566 | 2.752 | Gain |
| 7:19892333-24865416 | 2.323 | Loss |
| 1:39880100-44865111 | 2.249 | Gain |
| 21:14300000-48129895 | 1.697 | Loss |
| 6:44001018-48890019 | 1.337 | Gain |
| 14:68313344-73192868 | 1.254 | Gain |
| 16:28533080-33288593 | 1.113 | Gain |
| 4:107830618-112732008 | 1.018 | Gain |
| 14:14638574-19518098 | 0.978 | Loss |
| 2:203493353-208456605 | 0.977 | Loss |
| 2:89338546-94301797 | 0.879 | Loss |
| 7:4973084-9946166 | 0.759 | Loss |
| 4:132337576-137238967 | 0.729 | Gain |
| 6:146670058-151559059 | 0.631 | Gain |
| 2:29779516-34742767 | 0.593 | Loss |
| 4:1-4901391 | 0.515 | Gain |
| 3:108912337-113862897 | 0.406 | Loss |
| 20:43633053-48481169 | 0.376 | Loss |
| 2:183640343-188603595 | 0.343 | Gain |
| 16:66577187-71332699 | 0.331 | Loss |
| 22:27984309-32648360 | 0.327 | Loss |
| 10:87129481-91970006 | 0.317 | Loss |
| 4:44112526-49013916 | 0.275 | Gain |
| 3:1-87900000 | 0.250 | Gain |
| 1:69790174-74775186 | 0.209 | Loss |
| 7:54703916-59676998 | 0.204 | Loss |
| 1:149550373-154535385 | 0.202 | Loss |
| 1:84745212-89730223 | 0.201 | Loss |
| 4:171548710-176450100 | 0.194 | Gain |
| 2:129044566-134007817 | 0.191 | Loss |
| 9:111996860-116866287 | 0.180 | Loss |
| 10:14521581-19362106 | 0.165 | Loss |
| 7:84542415-89515497 | 0.152 | Loss |
| cx | 0.144 | - |

| | | |
|---|---|---|
| 19:54201568-59128983 | 0.139 | Loss |
| 1:199400497-204385509 | 0.138 | Gain |
| 2:54595778-59559030 | 0.137 | Gain |
| 18:19519313-24399140 | 0.136 | Gain |
| 7:74596249-79569331 | 0.126 | Gain |
| 2:59559031-64522282 | 0.118 | Loss |
| 5:176025659-180915260 | 0.111 | Loss |
| 5:127129643-132019243 | 0.109 | Loss |
| 18:63437765-68317592 | 0.103 | Gain |
| 6:102669041-107558042 | 0.100 | Gain |
| 11:53038275-57859935 | 0.097 | Gain |
| 2:69485536-74448787 | 0.096 | Gain |
| 12:114021985-118979462 | 0.094 | Gain |
| 1:79760199-84745211 | 0.089 | Loss |
| 10:29043161-33883686 | 0.084 | Gain |
| 4:186252885-191154276 | 0.080 | Gain |
| 1:44865112-49850124 | 0.076 | Gain |
| 9:14608286-19477714 | 0.073 | Loss |
| 1:169490423-174475434 | 0.055 | Loss |
| 1:24925063-29910074 | 0.052 | Loss |
| 16:52310647-57066159 | 0.051 | Loss |
| 1:144565361-149550372 | 0.048 | Gain |
| 5:136908846-141798447 | 0.042 | Loss |
| 10:125853694-130694220 | 0.037 | Gain |
| 2:84375293-89338545 | 0.033 | Gain |
| 4:73520876-78422267 | 0.026 | Gain |
| 17:38209511-42985699 | 0.023 | Loss |
| 10:67767374-72607900 | 0.019 | Gain |
| 7:1-4973083 | 0.013 | Gain |
| 15:20700000-102531392 | 0.008 | Gain |
| 7:9946167-14919249 | 0.005 | Gain |
| 6:9778004-14667005 | 0.005 | Loss |
| 5:171136057-176025658 | 0.005 | Gain |
| 4:29408351-34309741 | 0.004 | Loss |

**Supplementary Table 3 Model coefficients**

Genomic regions selected by the model as being predictive of progression with their associated coefficient of variation for the relative risk (i.e. $CV_{RR} = CV(e^{y\beta})$, where y is the matrix value) and status (e.g. relative gain or loss). Few are strong drivers of the model, and even in the top five coefficients are found in relatively few patients or samples. The ‘cx’ coefficient is the value added to the model to represent the overall complexity of the genome.

#### Supplementary Figures

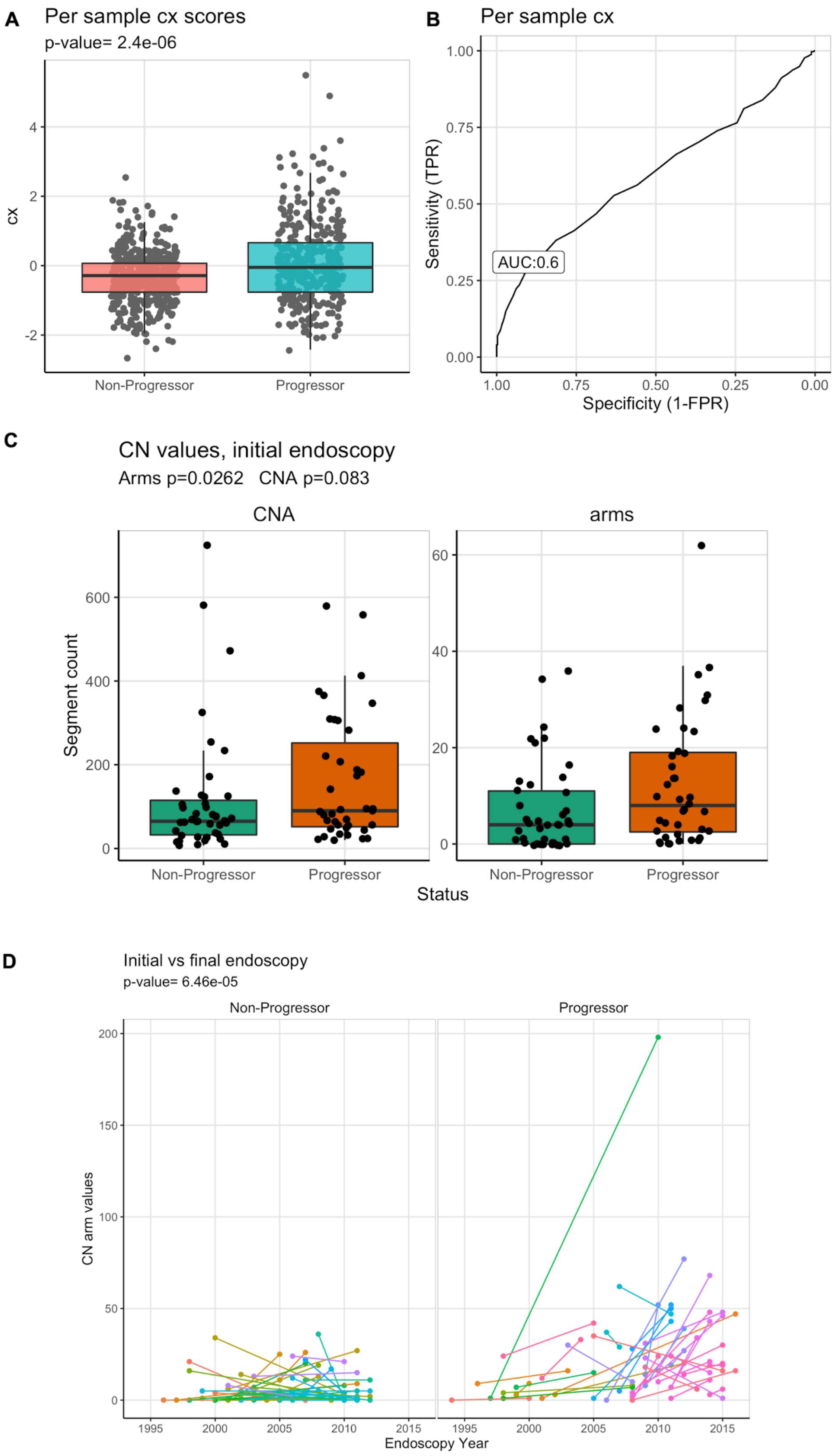


**Supplementary Figure 1:** (A) Per-sample variance in the genomic complexity (cx) values (y-axis) between progressors and non-progressors. While the difference is significant in a

Wilcoxon rank sum test, it only provides limited prognostic signal as the ROC curve in (B) shows. (C) Counts of segment values that are CN altered (y-axis) of the segments and arms split by progressors and non-progressors in the initial endoscopy. Chromosomal arms show a significant difference (p-value = 0.026, Wilcoxon rank sum test) in the number of CN alterations identified between the two groups. (D) Comparison of arm level CN values (y-axis) found at the initial vs the final endoscopies in progressors and non-progressors, showing significant changes (p-value 6.46x10-6, Wilcoxon rank sum test) occurring in the genomic landscape are apparent even in low-resolution WGS data.

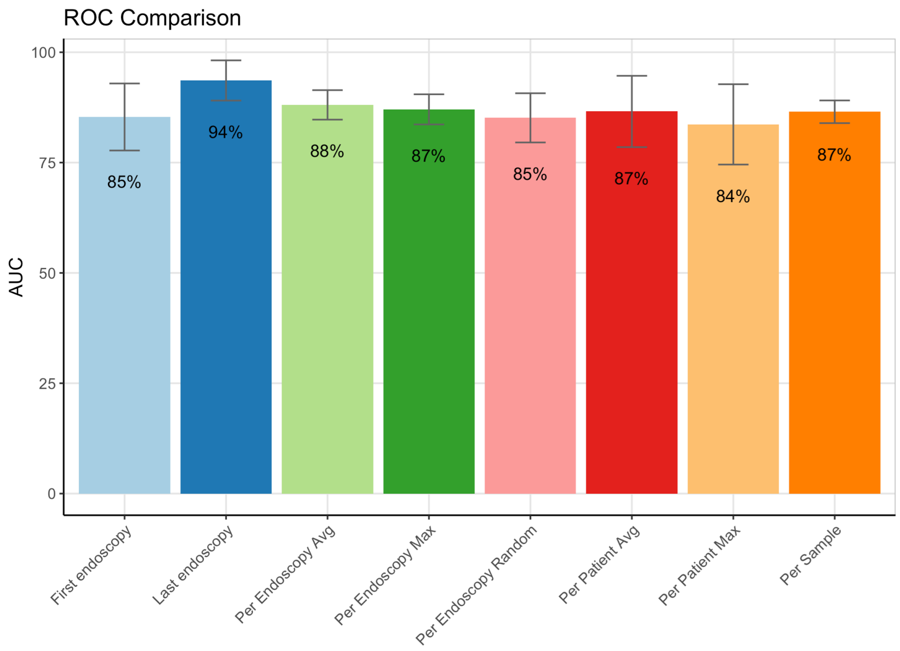


**Supplementary Figure 2:** ROC curves describing the accuracy of prediction for various aggregations of per-sample predictions. (A) predictions in samples in the first endoscopy, (B) predictions in samples in the last available endoscopy, (C) average prediction per endoscopy, (D) maximum prediction per endoscopy, (E) random sample per endoscopy, (F) average prediction per patient, excluding the final endoscopy, (G) maximum per patient prediction excluding the final endoscopy, and (H) the AUC from all 773 samples. (I) shows the AUC for A-G in a bar graph. Using the average or maximum prediction per endoscopy does not dramatically alter the AUC from the 0.87 per-sample rate, suggesting that a single sample may be sufficient.

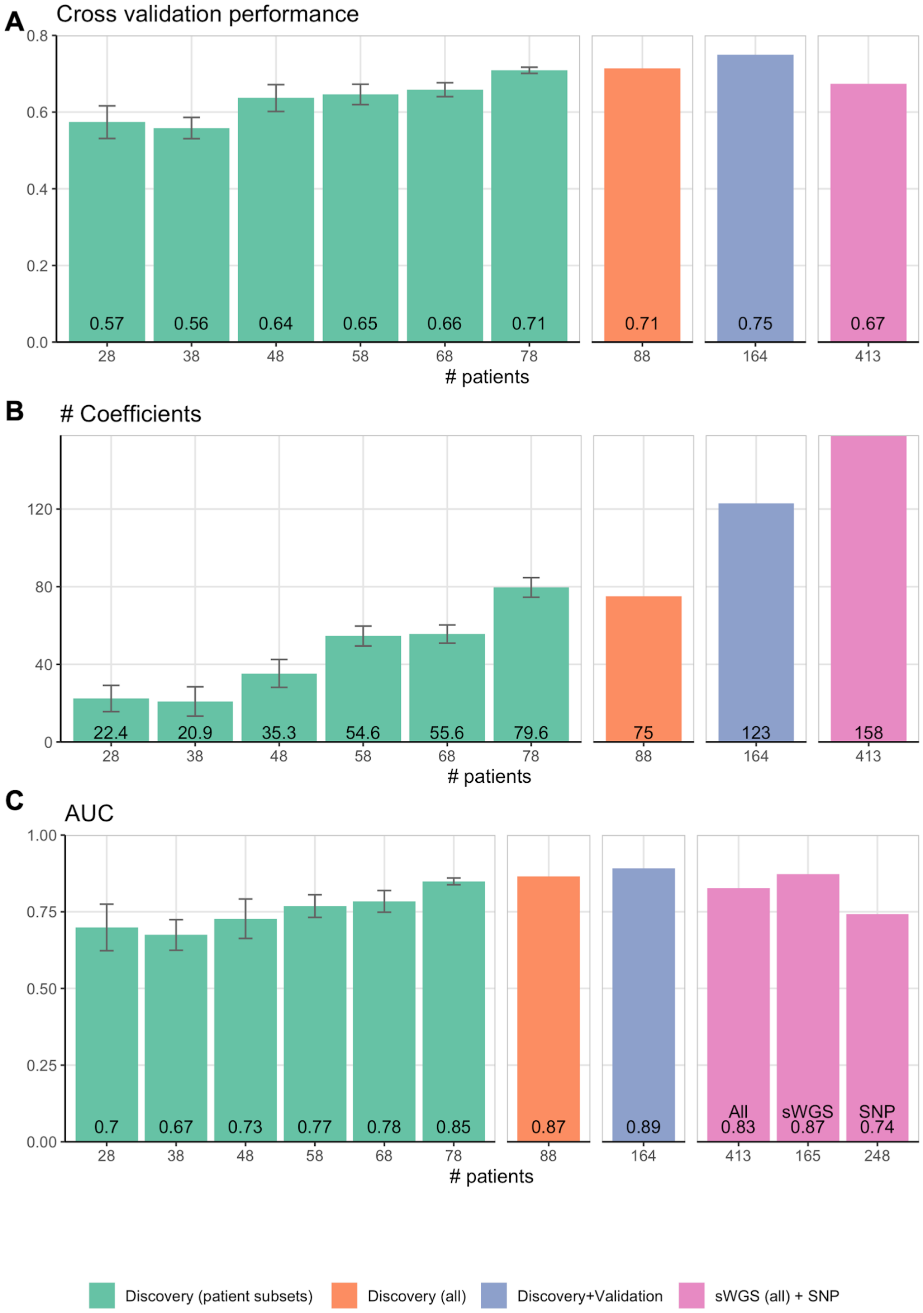


**Supplementary Figure 3:** Potential for improvement by training the model with increasing numbers of patients from the discovery cohort, combining the discovery and validation cohorts, and combining all sWGS (discovery and validation) data with the SNP data from the Seattle BE Study. In each model we assessed the (A) cross-validation performance, the (B) number of coefficients selected by the model, and finally the (C) AUC for a leave-one-out analysis. The green bars are all increasing numbers of patients used in training a model from the discovery cohort (repeated 10 times with randomly selected patients), the orange bar represents the full discovery cohort, the purple bar is the combined discovery (n=88) and validation (n=76) cohorts, and the pink bars are the combined sWGS and SNP patients (n=413).

The discovery and validation (all sWGS data) displays consistent improvement in accuracy (0.57 to 0.75) and AUC (0.7 to 0.89) as the number of patients increases. However, including the SNP data results in no improvement despite the increased number of patients indicating that the sWGS data alone provides more accurate prognostic information.

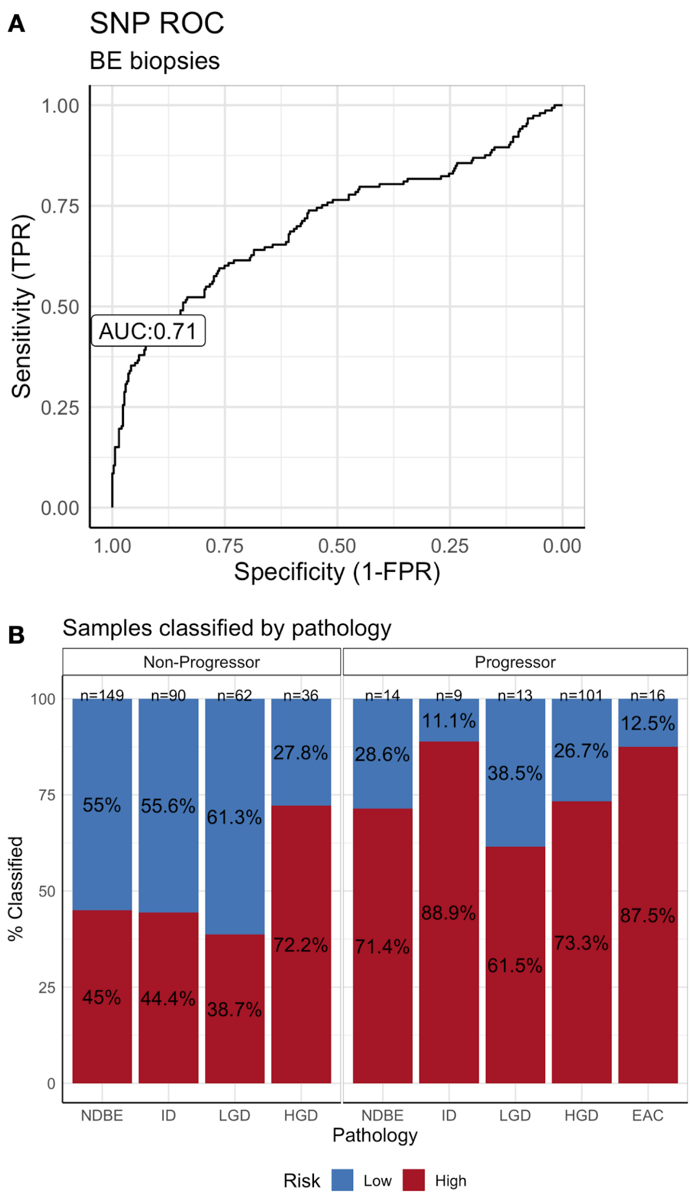


**Supplementary Figure 4:** Predicting the Seattle Barrett's Study SNP data using our sWGS model results in a lower AUC of 0.77 for all samples (including blood/gastric normals as NP controls) (A). Restricted to only Barrett's samples results in an AUC of 0.71 (B) Overall, the progressor samples show the same pattern of risk classification that sWGS samples do. The HGD group in the non-progressor patient group also indicates that our model would classify most of these as progressive.

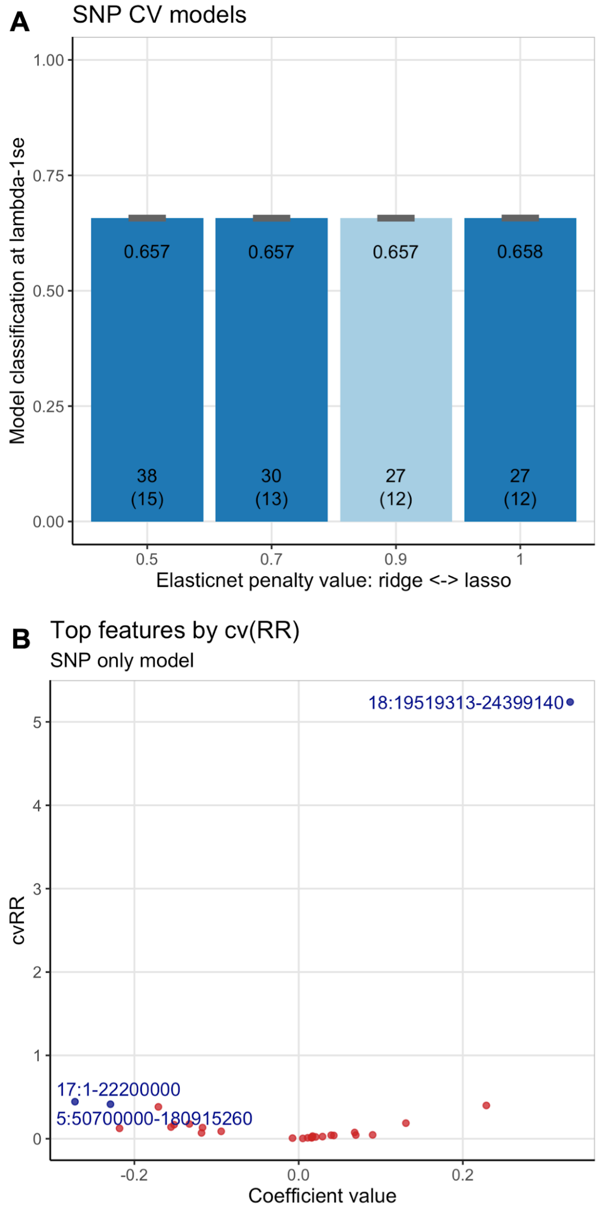


**Supplementary Figure 5:** (A) The cross-validation classification performance at each elastic net penalty value (penalty 0 had no non-zero coefficients), the light blue bar is the penalty value used in the sWGS model and is used for comparison. (B) Volcano plot for the cv(RR) value versus coefficient value for the 27 coefficients from the SNP data trained model. Compared to the coefficients from the sWGS model shown in Supplementary Fig. 4, the cv(RR) values (e.g. coefficient of variation for the relative risk, see Supplementary Table 3 for definition) are much lower.

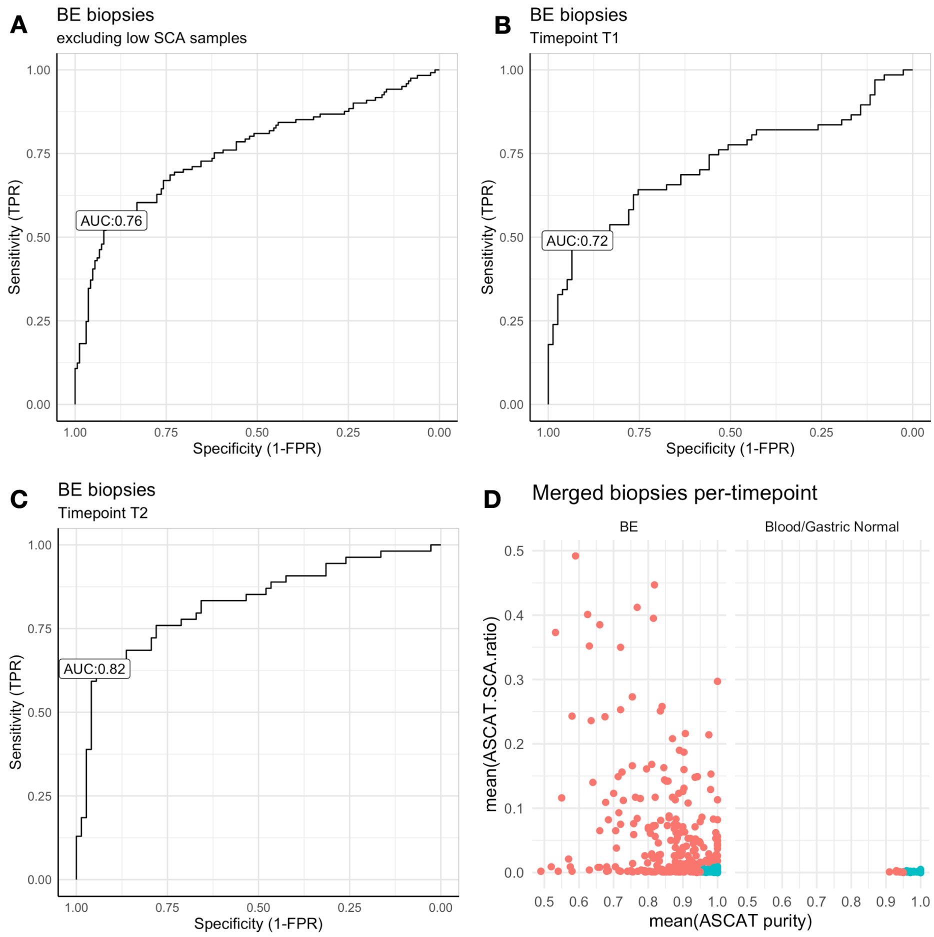


**Supplementary Figure 6**: (A) ROC curves for the SNP timepoint merged samples excluding those with less than 1% of the genome altered and the whole-genome duplicated non-progressor patients. (B) and (C) look at the same timepoint merged samples but only within the baseline (T1) and penultimate endoscopy (T2) groups respectively. Demonstrating that the model improves as the samples are taken nearer to EAC diagnosis. (D) Plots the mean ratio of the genome altered (y-axis) versus the computationally-derived purity value (x-axis) for all timepoint merged biopsies versus the blood/gastric normal samples. None of the normal samples have more than 1% of the genome altered, and all are >90% purity. Given the issues with assessing very pure mostly diploid samples, those samples in blue are excluded from the analyses in A-C.

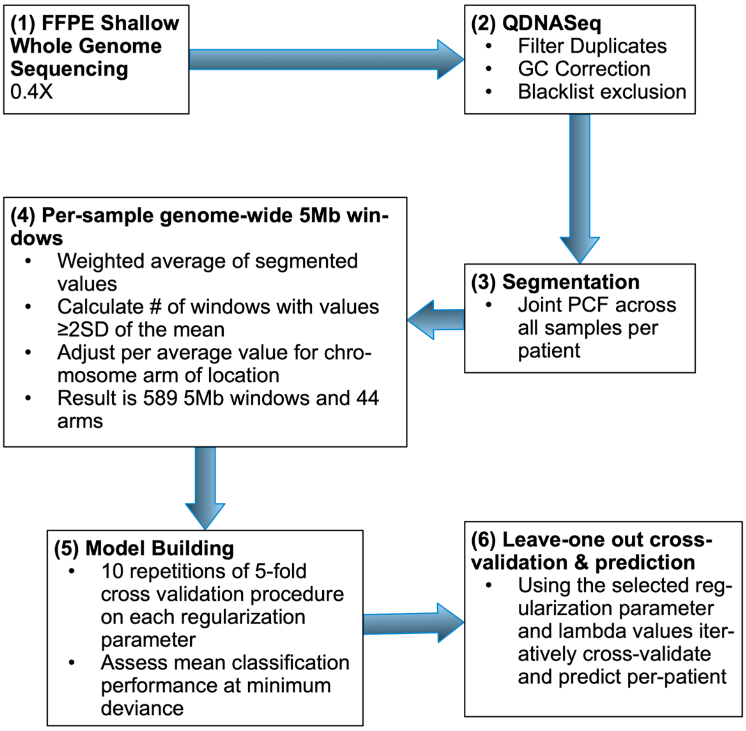


**Supplementary Figure 7:** The overall computational pipeline from generating the sWGS copy-number information to the and model training process and leave-one-out analysis.

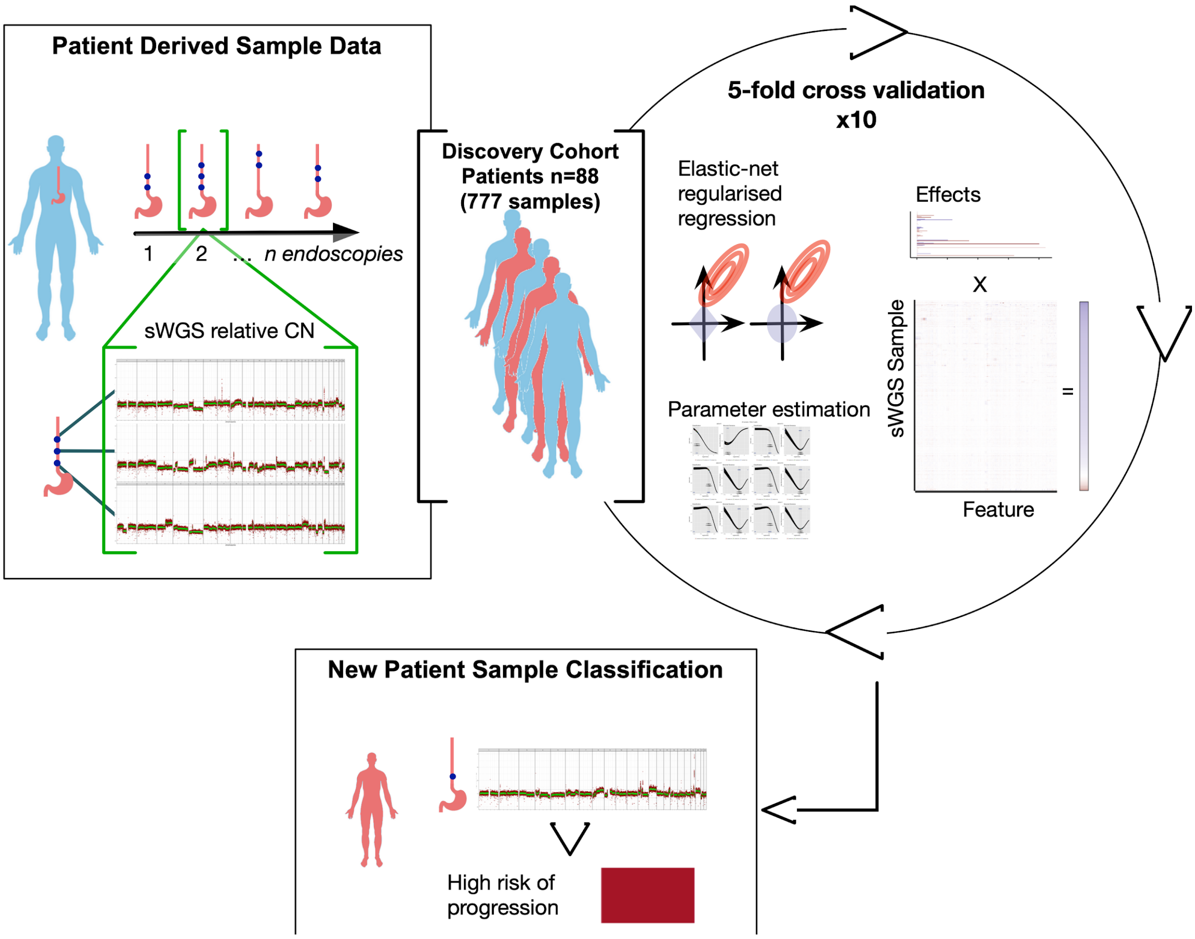


**Supplementary Figure 8:** Shows a brief overview of the model training process in our cohort. Single-quadrant samples from all patients over time are used in the cross-validation 80/20 training. New patient samples are segmented and windowed as described in Supp. Fig. 7 steps 3 and 4 then the risks are predicted and classified using the fully trained model.

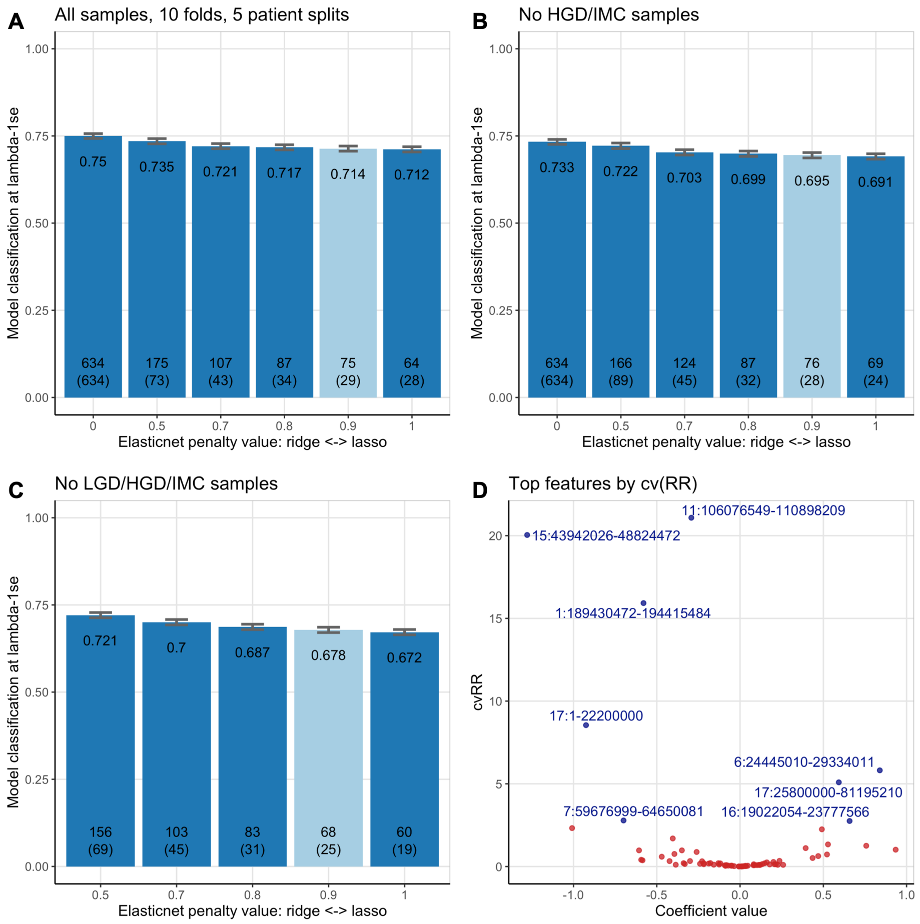


**Supplementary Figure 9:** Cross-validation model classification rates for different elastic net penalty values. The light blue bars show the penalty value used in the final model (0.9). Numbers at the bottom of the bars show the number of features selected in that model, the number in the parenthesis is the number of features that were stable across 75% or more of the folds. (A) shows the model classification for all samples in the discovery cohort, (B) excluding HGD or IMC, (C) excluding HGD, IMC and LGD. While the classification rate decreases it is still very similar to the all samples model. This indicates that the endpoint samples alone are not driving the model. (D) shows the 75 features with the top quartile by the coefficient of variation for the relative risk (cv(RR), see Supplementary Table 3) labeled.

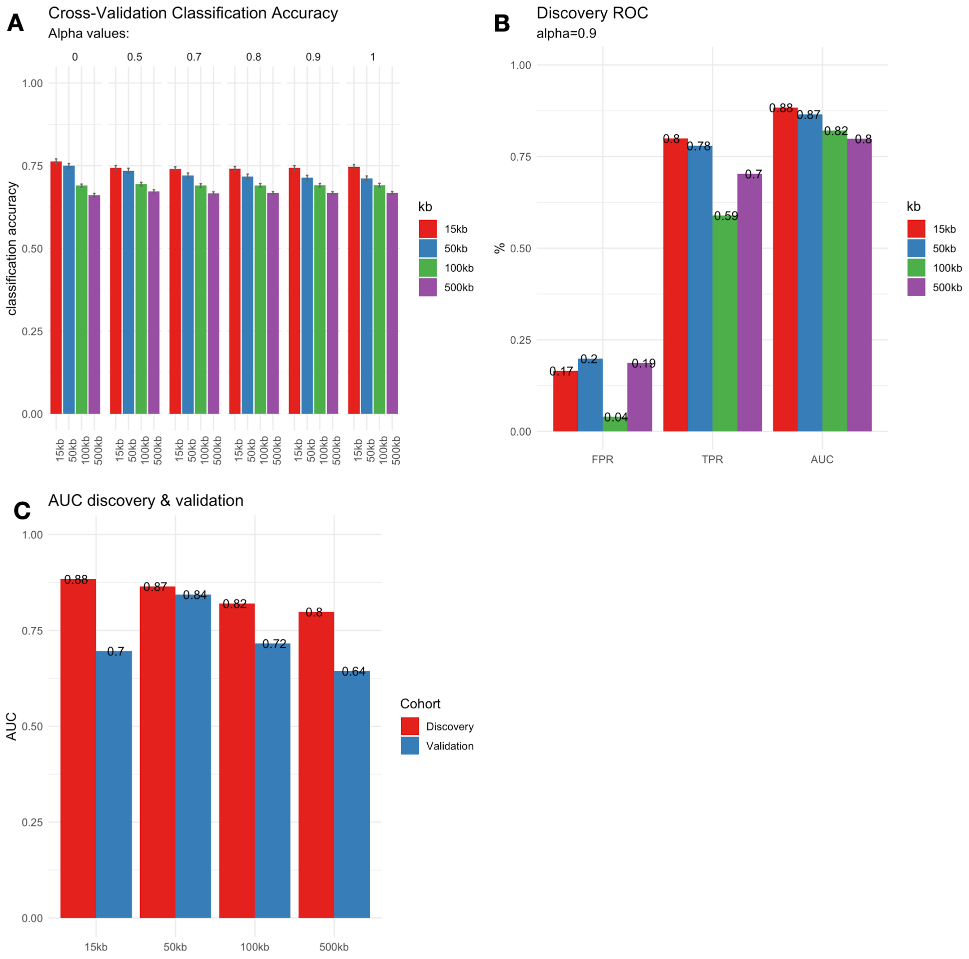


**Supplementary Figure 10:** (A) shows the cross-validation classification rate for each bin size (15kb, 50kb, 100kb, 500kb) at each elastic-net penalty value. The classification rate shows a consistent decline in order for each bin size. (B) Compares the AUCs for each bin size using leave-one out predictions for the discovery cohort at an elastic-net regression penalty of 0.9. Again 15kb shows the best AUC at 0.88, however 50kb is very highly concordant at 0.87. (C) shows the AUC comparison at each bin size for the leave-one out discovery cohort predictions versus the validation cohort model predictions. At 50kb the AUCs are 0.87 and 0.84 respectively while all other bin sizes show a much greater difference between the cohorts.

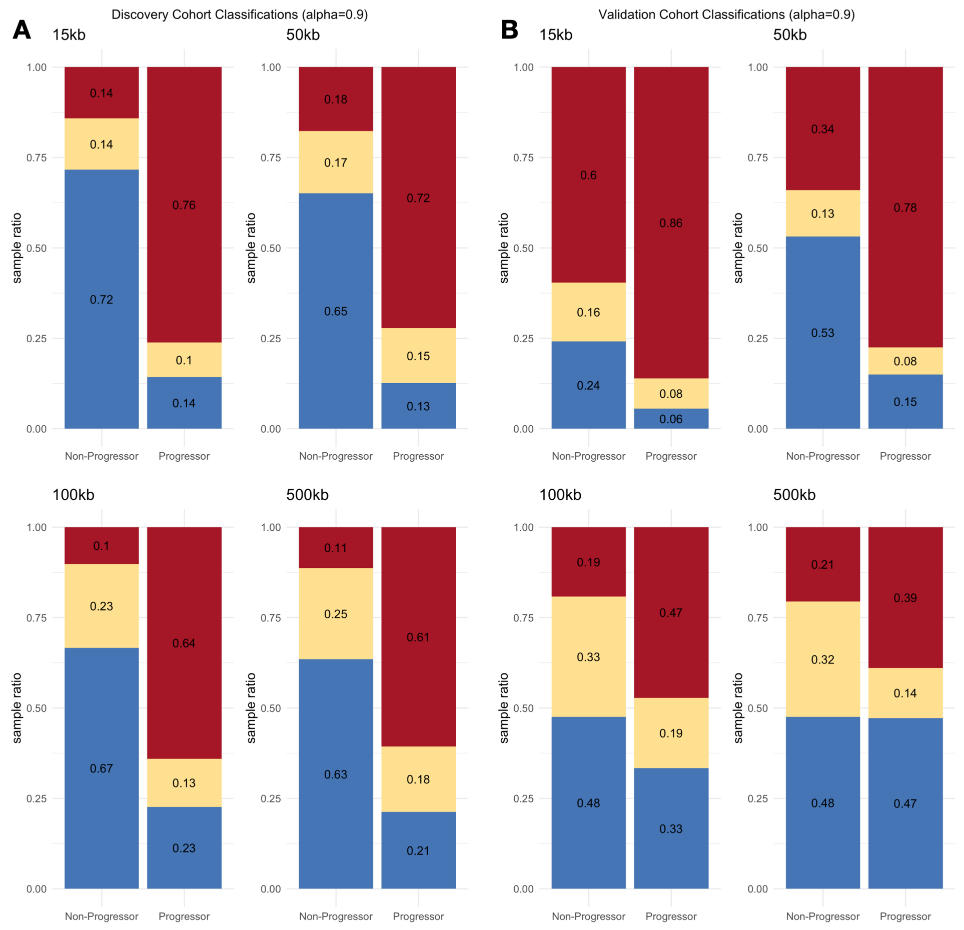


**Supplementary Figure 11:** Rate of sample classification by probability discretization per bin size for the discovery cohort leave-one-out predictions (A) and validation predictions (B). These confirm that 50kb is the best parameter to balance classification for type I and type II errors.

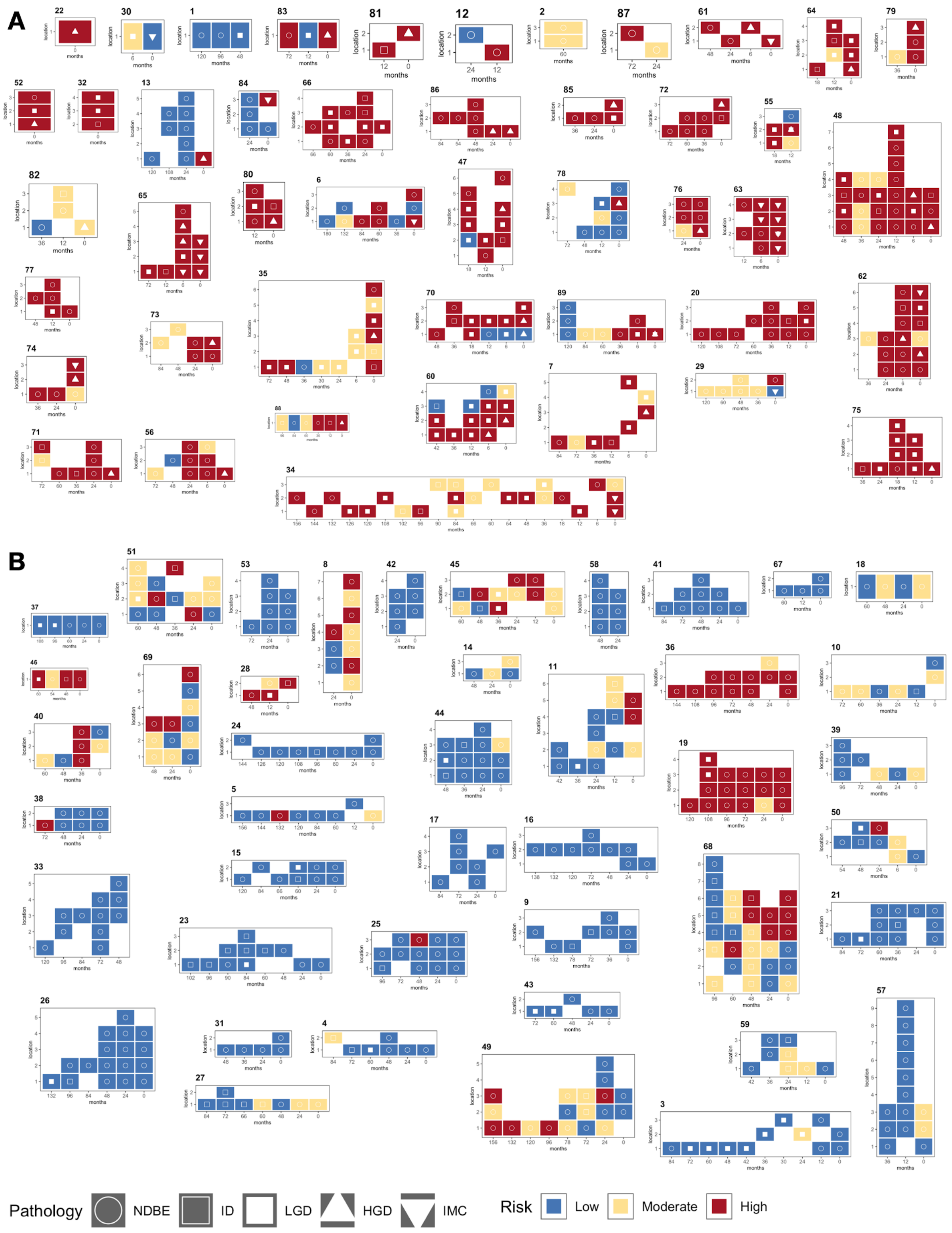


**Supplementary Figure 12**: Samples from the discovery cohort (n=773) for each progressor patient (n=45) in (A) plotted by the time until final endoscopy (x-axis) and esophageal location from the esophageal-gastric junction at the bottom to the length of the BE segment, or as many samples as were available for sequencing (y-axis). Each sample is colored by their risk class. Non-progressor patients are shown in (B) (n=43). These correspond to the mini heatmaps in Figure 2 with pathology per sample included.

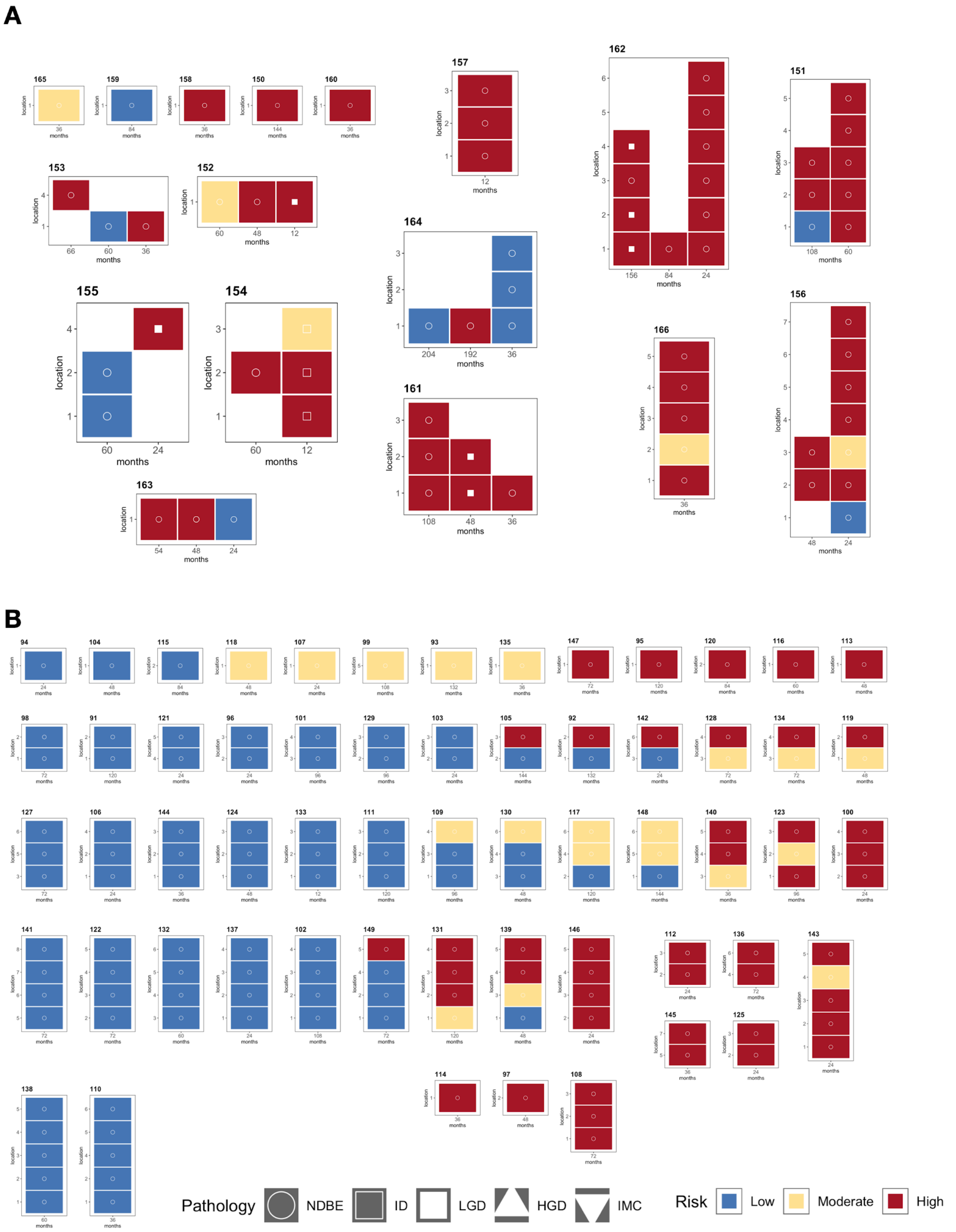


**Supplementary Figure 13**: Samples from the validation cohort (n=217) for each progressor patient (n=17) in (A) plotted by the time until final endoscopy (x-axis) and esophageal location from the esophageal-gastric junction at the bottom to the length of the BE segment, or as many samples as were available for sequencing (y-axis). Each sample is colored by their risk class. Non-progressor patients are shown in (B) (n=58).
